# Long-form hibernation-promoting factor coordinates 100S assembly and translational repression

**DOI:** 10.64898/2026.10.02.756224

**Authors:** Meng Wu, Yuanjie Bai, Changbin Zhang, Chuansen Yi, Rikuan Zheng, Lei Qi, Lingfan Zhu, Zhe Zhang, Chaomin Sun, Wenfei Li

## Abstract

Ribosome hibernation mediated by long-form hibernation-promoting factor (HPF) requires coordination of ribosome dimerization and translational repression, but how these functions are integrated remains unclear. Here, we present cryo-EM structures of native 100S disomes from a deep-sea *Bacillus* isolate, revealing a conformationally heterogeneous population of 100S disomes. Beyond the established roles of the HPF N-terminal domain (NTD) in decoding-center occlusion and the C-terminal domain (CTD) in dimerization, our analyses reveal additional contributions from the intervening Linker and a charged CTD surface positioned near the 16S rRNA 3′ terminus containing the anti-Shine-Dalgarno (aSD) sequence. Linker substitutions differentially affect translational repression while retaining the ability to form 100S disomes, with distinct effects of the two Linker segments. Mutation of the charged CTD surface enhances translational repression while retaining robust 100S formation, suggesting an additional role beyond CTD-mediated ribosome dimerization. Functional analyses in an *E. coli* cell-free translation system further show that the inhibitory activity of individual HPF regions varies with the translational context. Together, these findings support a modular model in which the NTD, Linker, and CTD make distinct but interconnected contributions to 100S assembly and translational repression.

## Introduction

Protein synthesis is a major energetic investment for bacterial cells and is tightly regulated in response to environmental fluctuations (Buttgereit & Brand, 1995; Wilson & Nierhaus, 2007; Russell & Cook, 1995; Szaflarski & Nierhaus, 2007). During prolonged nutrient limitation or other unfavorable conditions, bacteria reduce translational activity while preserving ribosomes for subsequent recovery when growth conditions improve (Wilson & Nierhaus, 2007; Starosta *et al*, 2014; Usachev *et al*, 2020b; Basu & Yap, 2016). Ribosome hibernation represents a conserved adaptive strategy in which bacteria reversibly suppress translation through formation of inactive ribosome complexes, including the 100S disome (Prossliner *et al*, 2018; Karari Njenga & Koch, 2026; Matzov *et al*, 2019; Ueta *et al*, 2013; Yoshida *et al*, 2004).

Specialized ribosome-associated factors mediate this process by stabilizing inactive ribosome conformations while preserving the potential for reactivation (Polikanov *et al*, 2012; Basu & Yap, 2017; Basu *et al*, 2020). In Gammaproteobacteria, such as *E. coli*, short-form hibernation-promoting factor (HPF) cooperates with ribosome modulation factor (RMF) to promote 70S dimerization and inhibit translation through coordinated interactions at the decoding center and the 16S rRNA 3′ aSD region (Polikanov *et al*, 2012; Kato *et al*, 2010; Ueta *et al*, 2008; Beckert *et al*, 2018; Prossliner *et al*, 2021; Yoshida *et al*, 2002). In contrast, many Firmicutes utilize a single long-form HPF protein containing an N-terminal domain (NTD), a flexible linker region, and a C-terminal domain (CTD) (Khusainov *et al*, 2017; Beckert *et al*, 2017; Franken *et al*, 2017; Mishra *et al*, 2018; Flygaard *et al*, 2018). Structural studies of long-form HPF-containing 100S particles have revealed that the NTD occupies the decoding center, whereas the CTD mediates intermolecular interactions between ribosomes to promote disome formation (Beckert *et al*, 2017; Matzov *et al*, 2017; Franken *et al*, 2017; Basu *et al*, 2018; Khusainov *et al*, 2017; Gohara & Yap, 2018; Flygaard *et al*, 2018).

Recent cryo-EM studies further suggest that hibernating ribosomes can adopt different conformational states rather than a single inactive configuration, highlighting structural diversity within hibernating ribosome populations (Khusainov *et al*, 2017; Beckert *et al*, 2018; Matzov *et al*, 2017, 2019; Gohara & Yap, 2018). Together, these studies have provided a framework for understanding HPF-mediated ribosome hibernation, while the organization of conformational states in native 100S populations and the functional coordination among different HPF regions remain to be further defined.

Here, using native 100S disomes isolated from the deep-sea *Bacillus* sp. DY0427 (hereafter *Bacillus*), we combine cryo-EM with biochemical and functional analyses to define how long-form HPF coordinates ribosome dimerization and translational repression. Our structure reveals a conformationally heterogeneous 100S population containing a predominant non-rotated/non-rotated state and a distinct non-rotated/rotated state. Through systematic dissection of HPF domains, we show that the NTD, Linker, and CTD contribute distinct but interconnected functions, including decoding-center inhibition, ribosome dimerization, and regulation of the aSD-containing 16S rRNA 3 ′ terminus. Together, these findings reveal how long-form HPF integrates multiple functional modules to coordinate 100S assembly and translational repression.

## Results

### Conformational heterogeneity of the 100S disome

To investigate ribosome hibernation in a deep-sea bacterium, we cultivated *Bacillus* — an anaerobic isolate from a deep-sea hydrothermal vent, and harvested cells at late stationary phase, when 100S ribosome accumulation is commonly observed. Cell lysates were subjected to sucrose gradient centrifugation (SGC) to separate native 100S hibernating disomes from 70S monomers (Fig. 1A). Fractions corresponding to the 100S disome peak were collected and prepared for cryo-EM, yielding a 3D reconstruction at an overall resolution of 3.7 Å (Fig. 1A, Fig. S1, Table S1).

**Figure 1.**
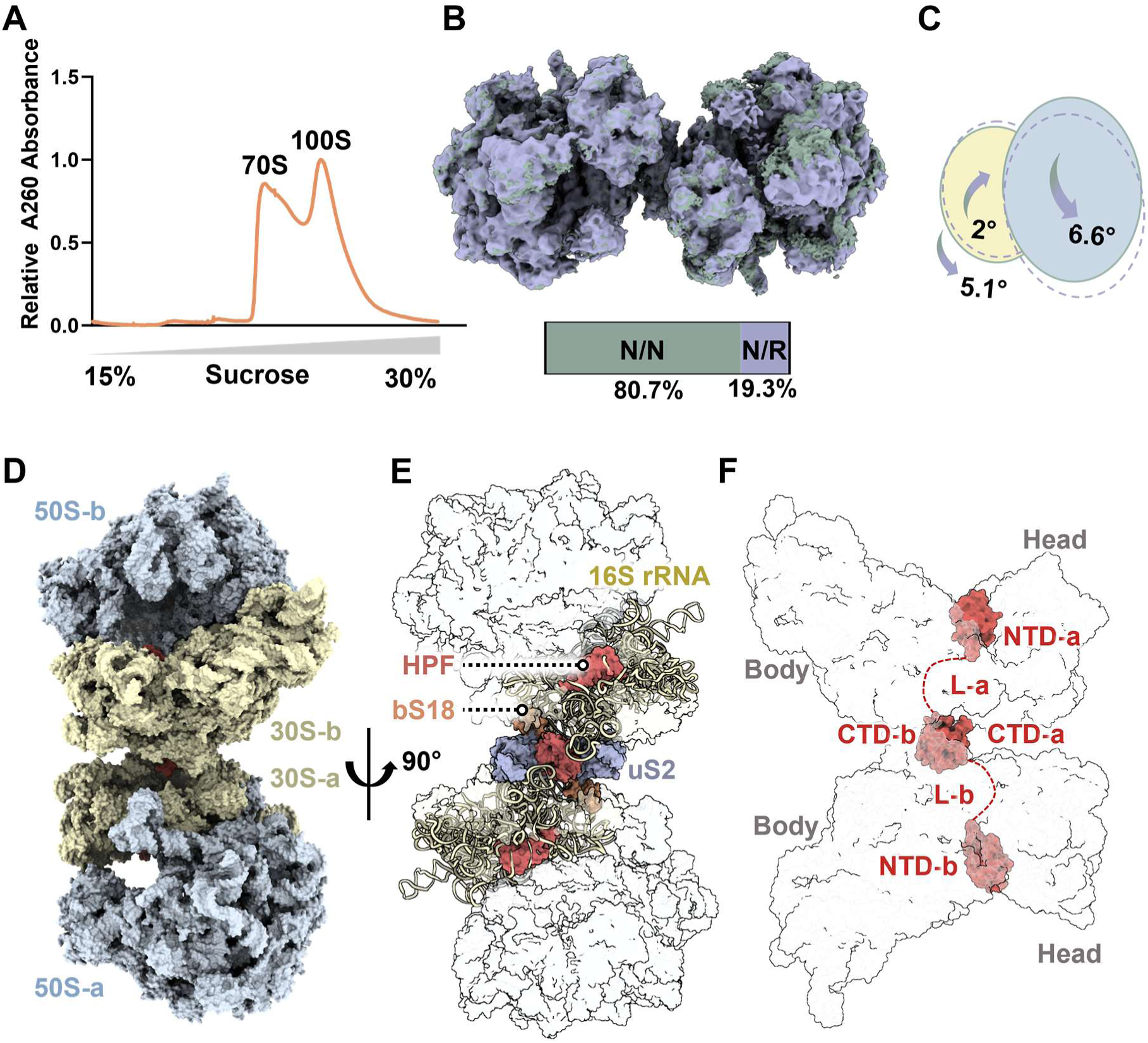
Structural architecture and conformational heterogeneity of the 100S disome. **(A)** Sucrose density gradient profile of native ribosomes from *Bacillus*, showing the 70S and 100S fractions used for cryo-EM analysis. **(B)** Cryo-EM density maps revealing conformational heterogeneity of the 100S disome with the non-rotated/non-rotated (N/N) state in green and the non-rotated/rotated (N/R) state in purple. The bar chart at the bottom displays the relative proportions of the N/N (80.7%) and N/R (19.3%) states. (**C**) Schematic representation of inter-subunit and inter-ribosomal rotational angles defining the transition between the N/N and N/R states, illustrating a 5.1° rotation of the rotated monomer relative to the non-rotated monomer, alongside dynamic subunit LSU/SSU movements (6.6° and 2.0°). **(D)** Overall cryo-EM density map of the *Bacillus* 100S disome in surface representation, showing the top-to-bottom arrangement (50S-b, 30S-b, 30S-a, and 50S-a). (**E**) Cartoon representation of the 100S disome interface turned 90° relative to (**D**). The long-form HPF molecules (red) and key surrounding 30S structural components—including 16S rRNA (tan), ribosomal proteins uS2 (purple), and bS18 (orange)—are highlighted. (**F**) Schematic outline depicting the modular domain organization and binding trajectory of long-form HPF monomers (HPF-a and HPF-b) spanning the 30S decoding center (NTD) through the intervening flexible linkers (L-a, L-b) to the central dimerization interface (CTD-a, CTD-b) between the two ribosomal 30S bodies.

The 100S reconstruction reveals an approximately symmetric arrangement of two 70S ribosome monomers, with the 50S subunits positioned on the outer sides of the complex and the two 30S subunits facing each other at the inter-ribosomal interface (Fig. 1B). Extensive 3D classification of the disome dataset uncovered conformational heterogeneity within the hibernating population (Fig. 1B-C, Fig. S1). The majority of particles (80.7%) adopted a symmetrical non-rotated/non-rotated (N/N) state, with both 70S ribosome monomers in an unratcheted conformation (Fig. 1B).

A distinct subpopulation (19.3%) adopted an asymmetrical non-rotated/rotated (N/R) disome state in which the two monomers adopt non-rotated and rotated states, respectively (Fig. 1B). Structural superposition of the N/N and N/R disome models reveals an overall rotation of approximately 5.1° between the two states (Fig. 1C). Alignment of the individual subunits reveals that the small subunit (SSU) exhibits a rotation of approximately 2°, while the large subunit (LSU) undergoes a rotation of approximately 6.6° (Fig. 1C).

Sequence alignment showed that *Bacillus* HPF contains conserved N- and C-terminal regions characteristic of long-form HPFs, with a distinct Linker region (Fig. S2). Structural analysis reveals that HPF occupies the interface between the two small subunits, while each ribosome monomer maintaining a canonical 70S architecture (Fig. 1D-E). Within each 70S monomer of the 100S disome, full-length HPF exhibits a clear modular domain architecture: an N-terminal domain (HPF^NTD^), an intervening flexible linker (HPF^L^), and a C-terminal domain (HPF^CTD^), that spans the inter-ribosomal interface (Fig. 1F). The NTD, CTD, and the associated ribosomal proteins uS2 and bS18 show well-resolved local density, supporting structural interpretation of these regions (Fig. S3). Superposition of the HPF models shows that the HPF-binding configuration is maintained across the two conformational states despite the rotational rearrangement of one ribosome monomer (Fig. S2B). The co-existence of the N/N and N/R populations indicates that 100S disomes can adopt distinct rotational conformations within the hibernating ensemble. This conformational heterogeneity indicates that the hibernating ribosome can adopt multiple structural states rather than a single inactive configuration.

### Steric blockade of tRNA and mRNA binding sites by HPF^NTD^

HPF^NTD^ occupies the decoding center on the 30S subunit, adopting a βαβββα fold (β1-β4 and α1-α2) (Fig. 2A). Structural analysis reveals that HPF^NTD^ wedges into the core of the decoding center, where it establishes extensive contacts with key ribosomal framework elements, including helix 44 (h44) and helix 18 (h18) of the 16S rRNA, helix 69 (H69) of the 23S rRNA, and the ribosomal protein uS12 (Fig. 2B).

**Figure 2.**
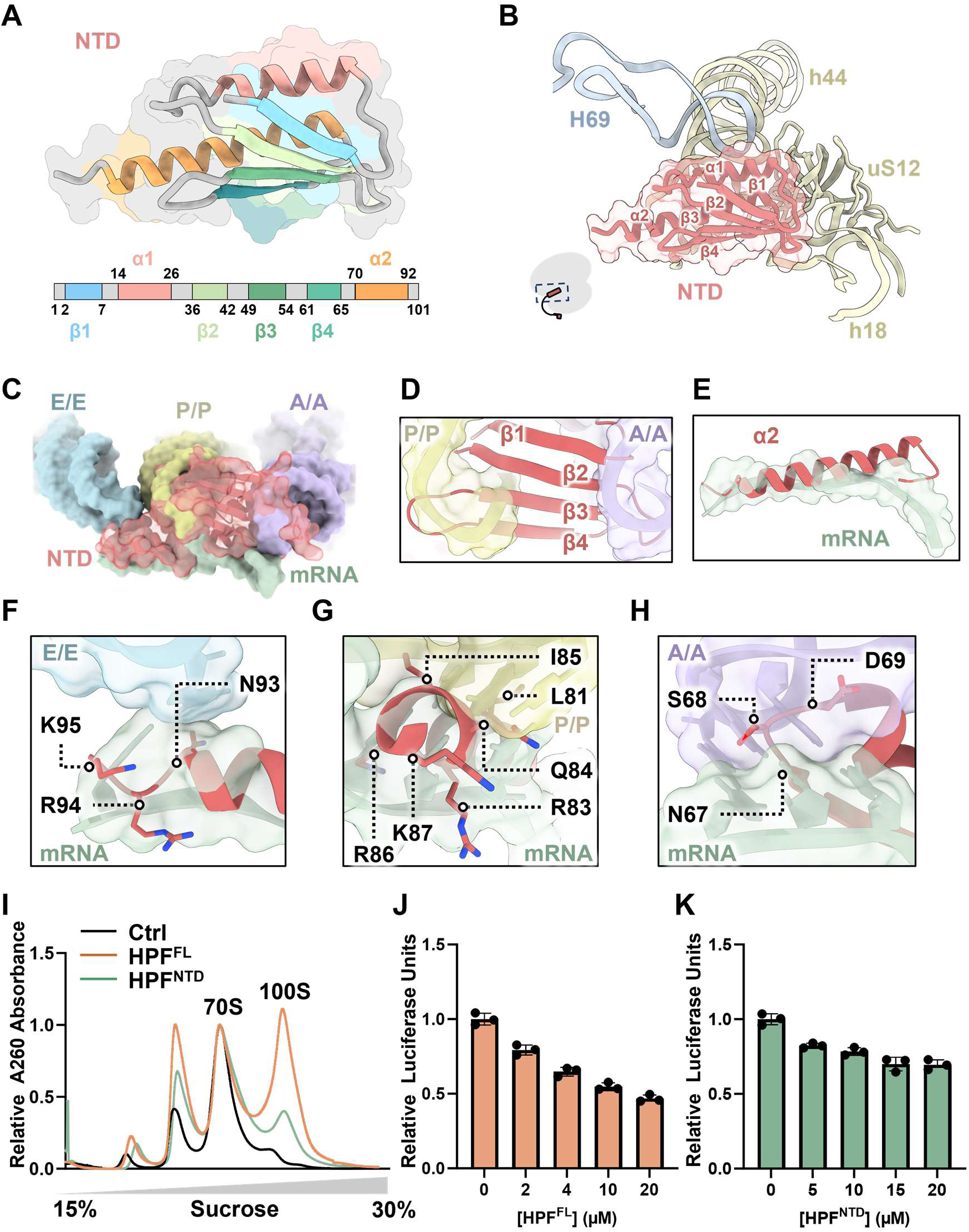
Structural and functional analysis of HPF^NTD^. **(A)** Structure topology of HPF^NTD^, showing α-helices (α1 and α2) and four β-strands (β1-β4), colored according to residue positions**. (B)** Close-up view of HPF^NTD^ (red) inserted into the decoding center. The inset indicates the global viewing angle relative to the 70S ribosome. **(C)** Overview of the spatial relationship between HPF^NTD^ and the tRNA-binding sites at the decoding center. **(D)** Superposition of HPF^NTD^ with A/A- and P/P-site tRNAs, showing the spatial overlap between the β1-β4 of the NTD and the tRNA-binding regions. (**E**) Close-up view of the α2 helix positioned within the mRNA channel. (**F-H**) Close-up views of representative interactions between NTD and the mRNA path in E, P and A sites, respectively. Residues involved in the indicated interactions are labeled. (**I**) Sucrose gradient profiles of 70S ribosomes (∼2 µM) incubated without HPF or with HPF^FL^ or HPF^NTD^ at a 1:1 molar ratio. **(J-K)** IVT assays showing the effects of HPF^FL^ and HPF^NTD^ using Nanoluciferase (NLuc) reporter in the *Bacillus* cell-free system. Data are presented as mean ± s.d. (n = 3).

HPF^NTD^ establishes a multi-site steric blockade within the functional centers of the ribosome (Fig. 2C). Specifically, the central β-sheet (β1-β4) overlaps with the canonical binding spaces for A-site and P-site tRNAs (Fig. 2D). Simultaneously, helix α2 projects directly into the mRNA entry channel (Fig. 2E). Detailed contact analysis shows that specific basic and polar residues within HPF^NTD^ are positioned along the mRNA path across the E-, P-, and A-site regions (Fig. 2F-H). Structural superposition reveals that the overall three-dimensional fold of HPF^NTD^ is highly conserved across representative long-form and short-form HPF factors from diverse bacterial lineages, with low RMSD values (0.94-1.20 Å) (Fig. S4).

To determine whether this conserved structural module is sufficient to support HPF-mediated disome assembly and translational repression, we next examined the ability of isolated HPF^NTD^ to promote 100S assembly and inhibit translation. Sucrose gradient analysis showed that the isolated HPF^NTD^ did not efficiently promote 100S disome formation compared to the HPF^FL^ (Fig. 2I, Fig. S5). This result is consistent with the structural organization of HPF, as the NTD occupies the decoding center whereas the CTD provides the inter-ribosomal interface required for disome assembly. *In vitro* translation (IVT) assays using a NanoLuciferase reporter system in our established *Bacillus* cell-free translation system showed that that HPF^NTD^ was substantially less inhibitory than full-length HPF, retaining approximately 70% of relative NLuc expression even at 20 μM (Fig. 2J and K). Thus, despite its conserved positioning within the decoding center, isolated HPF^NTD^ alone exhibits only limited translational inhibitory activity under the dynamic conditions of protein synthesis.

Together, these results indicate that the structural positioning of HPF^NTD^ within the decoding center can sterically interfere with tRNA and mRNA binding, while efficient 100S assembly and strong translational repression require additional regions of HPF.

### HPF^CTD^ mediates ribosome dimerization and modulates translational repression

Dimerization of two 70S monomers into the 100S disome is mediated by interactions between HPF^CTD^s (Fig. 3A). Each HPF^CTD^ adopts a four-stranded antiparallel β-sheet module (β5-β8) accompanied by helix α3 (Fig. 3A). The two CTDs form a clasp-like dimer, in which β5 from one CTD extends toward β8 of the opposing CTD, while the central β6-β8 sheets form a compact hydrophobic interface (Fig. 3B,C, Fig. S6A). Structural comparison with the crystal structure of isolated *Staphylococcus aureus* HPF^CTD^ (Usachev *et al*, 2020a) confirms that this CTD fold and dimerization interface are conserved (RMSD = 0.81 Å) (Fig. S6B).

**Figure 3.**
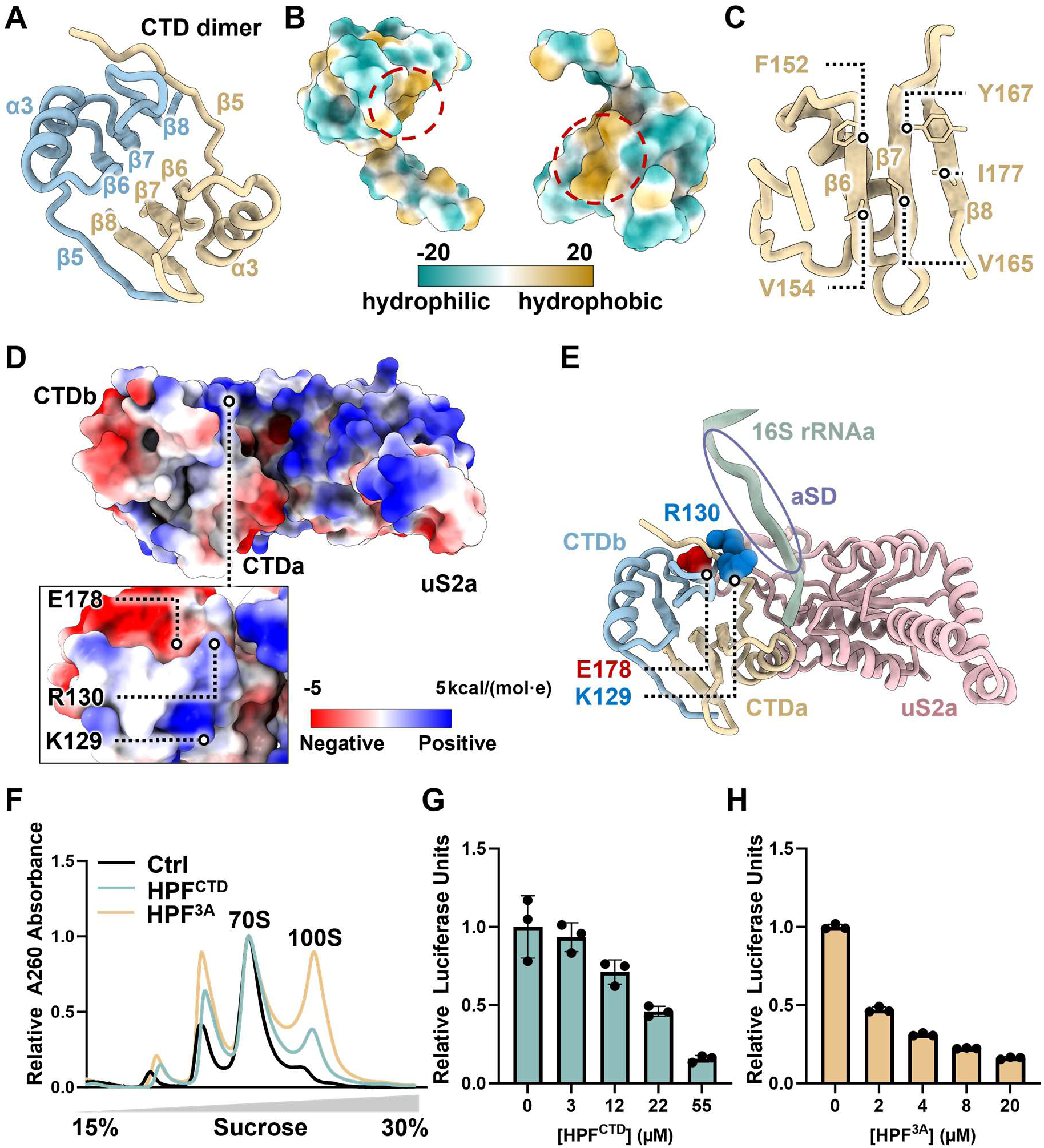
Structural analysis of the HPF^CTD^ dimerization interface and its inhibitory role in ribosome assembly and translation. **(A)** Overall representation of the HPF^CTD^ dimer in ribbon format, showing secondary structure elements. **(B)** Hydrophobicity surface representation of the two HPF^CTD^ monomers. The red dashed circles indicate the hydrophobic interface mediating dimerization. **(C)** Close-up view of the central face-to-face β-sheets interface formed by β6-8, with key hydrophobic residues (F152, V154, V165, Y167, I177) shown as sticks. **(D)** Electrostatic surface representation of the HPF^CTD^ and adjacent uS2. The lower-left inset highlights the positively charged surface pocket containing residues Lys129, Arg130, and Glu178. **(E)** Model showing the trajectory of the 16S rRNA 3′ terminus (containing the anti-SD sequence) positioned adjacent to the electrostatic pocket. Key residues K129, R130 (monomer a), and E178 (monomer b) are shown as spheres. **(F)** Sucrose gradient profiles of 70S ribosomes incubated in the absence of HPF (Ctrl, black), or with HPF^CTD^ (cyan), or with the triple mutant HPF^3A^ (orange) at a 1:1 molar ratio. **(G-H)** IVT assays showing the concentration-dependent translational repression of HPF^CTD^ and HPF^3A^ using the NLuc reporter in the *Bacillus* cell-free system. Data are presented as mean ± s.d. (n = 3).

In addition to the inter-ribosomal contact surface, the CTD presents an electrostatic surface facing the 30S subunit, positioned adjacent to the flexible 16S rRNA 3′ terminus containing the aSD sequence, which extends from helix 44 (Fig. 3D-E, Fig. S6C). To examine the functional role of this surface, we generated a triple mutant (HPF^3A^: K129A/R130A/E178A), targeting three charged residues located on this electrostatic CTD surface adjacent to the 16S rRNA 3′ terminus (Fig. 3D-E).

To examine the contribution of the CTD region to disome assembly, we first evaluated isolated HPF^CTD^ and full-length HPF^3A^ by sucrose gradient analysis (Fig. 3F, Fig. S5). Isolated HPF^CTD^ showed markedly reduced 100S formation efficiency compared to full-length HPF (Figs.2I and 3F). This indicates that, while the CTD provides the physical dimerization interface, the isolated domain alone is insufficient for robust 100S assembly, requiring the full-length multi-domain framework. In contrast, HPF^3A^ maintained robust 100S assembly activity, displaying an efficiency comparable to HPF^FL^ (Figs. 2I and 3F).

We next evaluated the effect of this CTD surface on translational repression using IVT assays. The isolated CTD exhibited minimal inhibitory activity, consistent with the absence of the decoding-center inhibitory activity provided by the NTD (Fig. 3G). In contrast, HPF^3A^ produced stronger translational repression than wild-type HPF across corresponding concentrations (Figs. 2J and 3H). Together, these results indicate that mutation of the charged CTD surface enhances translational repression while retaining robust 100S assembly.

### The HPF linker differentially regulates ribosome assembly and translational repression

The disome reconstruction further revealed the spatial position of the HPF Linker-CTD region relative to the 16S rRNA 3′ terminus. The 16S rRNA 3′ tail, containing the conserved aSD sequence, extends from helix 44 toward a region flanked by the Linker-CTD junction and ribosomal proteins (Figs. S6C and S7A). Although local structural flexibility precludes definitive assignment of individual side-chain interactions between the Linker and the aSD region, continuous backbone density is observed along the 16S rRNA 3′ tail (Fig. 4A).

**Figure 4.**
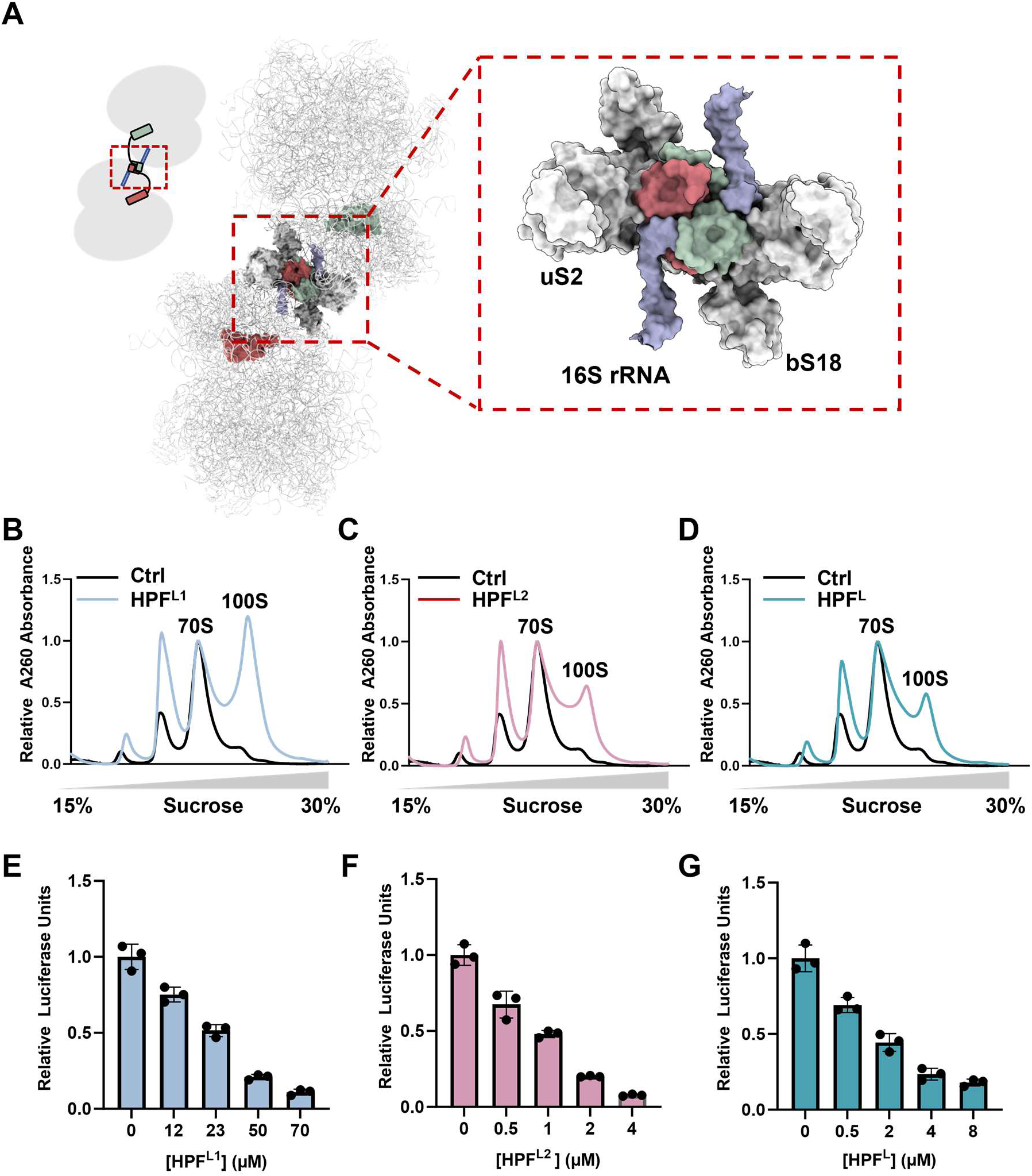
Structural and functional analysis of the HPF linker-CTD region near the 16S rRNA 3′ terminus. **(A)** Structural organization of the HPF Linker-CTD region adjacent to the 16S rRNA 3′ extension (containing the anti-SD sequence) and ribosomal proteins uS2 and bS18. The left insets depict the overall viewing angle relative to the 100S disome. The close-up view highlights the spatial proximity of the two HPF monomers (colored red and green, respectively) to the 16S rRNA 3′ tail (purple/dark blue) and surrounding proteins uS2 and bS18. **(B-D)** Sucrose gradient profiles of 70S ribosomes incubated with HPF^L1^, HPF^L2^ and HPF^L^ at a 1:1 molar ratio, compared with the control without protein. (**E-G**) IVT assays showing the concentration-dependent translational repression of HPF^L1,^ HPF^L2^ and HPF^L^ using the NLuc reporter in the *Bacillus* cell-free system. Data are presented as mean ± s.d. (n = 3).

To understand how the HPF Linker is positioned relative to hibernation factors in other bacterial systems, we compared our structure with hibernating ribosome complexes from other bacterial lineages (Fig. S7). In *E. coli*, which expresses short-form HPF (*e*HPF) lacking a CTD, the aSD sequence within the 16S rRNA tail is engaged by the separate RMF (Fig. S7B) (Beckert *et al*, 2018). In *Mycobacterium tuberculosis*, the long-form factor RafH is positioned near the 16S rRNA 3′ region without promoting dimerization, and residues within its Linker region have been shown to interact with nucleotides in the 16S rRNA aSD region (Fig. S7C) (Kumar *et al*, 2024). These comparisons suggest that, although different bacterial factors vary in sequences and disome assembly capacity, the region adjacent to the 16S rRNA 3′ terminus is recurrently occupied or engaged by hibernation factors (fig. S8).

To dissect the functional contributions of the intervening linker (residues 102-123), we generated flexible Gly-Ser (GS) substitution variants targeting individual Linker segments (Figs. S2 and S5). The 22-residue Linker was divided at residue 113 into Linker 1 (residues 102-112, IKTNFILPEAV) and Linker 2 (residues 113-123, AVLDQVEEDEL) (Fig. S2). Linker 1 directly connects the NTD (α2) with the downstream region and may influence the functional state of the NTD-containing region, whereas Linker 2 contains an acidic cluster (D116 and E120-E123) immediately preceding the HPF^CTD^ (β5), in a region positioned near the 16S rRNA 3′ terminus. We therefore generated three GS substitution variants, including HPF^L1^, HPF^L2^, and HPF^L^, in which Linker 1, Linker 2, or the entire Linker was replaced by GS sequences, respectively (Fig. S5).

Sucrose gradient analysis revealed that all three Linker replacement variants retained the ability to promote 100S disome formation in the presence of purified 70S ribosomes (Fig. 4B-D). HPF^L1^ exhibited the highest 100S assembly efficiency, whereas HPF^L2^ and HPF^L^ showed moderately reduced disome formation (Fig. 4B-D). To determine whether these differences reflected altered basal ribosome association, we performed sucrose cushion co-pelleting assays using FLAG-tagged HPF variants incubated with purified 70S ribosomes. Western blot analysis showed comparable ribosome association for all Linker variants (Fig. S7D).

We next examined the impact of Linker replacement on translational repression using the *Bacillus in vitro* translation assays (Fig. 4E-G). The variants displayed distinct inhibitory activities, with HPF^L2^ producing stronger repression than HPF^L^, whereas HPF^L1^ showed markedly weaker inhibition (Fig. 4E-G). HPF^L1^ required substantially higher protein concentrations to achieve comparable levels of repression, whereas HPF^L2^ consistently produced stronger inhibition than HPF^L^ across the tested concentrations (Fig. 4E-G). These results indicate that the two Linker segments differentially modulate HPF activity. The weaker inhibitory activity of HPF^L1^ suggests that alterations to Linker 1 can affect the functional activity of the NTD-containing region, whereas the retention of substantial inhibitory activity in HPF^L2^ indicates that Linker 2 is not strictly required for translational repression. Together, these findings identify the HPF Linker as a modulatory element that differentially tunes100S assembly and translational repression.

### A modular HPF integrates ribosome dimerization and translational repression

Structural comparisons of hibernating ribosome complexes showed that the NTD is conserved between the long-form HPF analyzed here and the short-form HPF of *E. coli* (Fig. 5A,B, Fig. S4D). Long-form HPF factors, such as those from *Bacillus* and *Thermus thermophilus*, contain a CTD that contributes to inter-ribosomal association and 100S disome formation. In contrast, *E. coli* uses a short-form HPF lacking the CTD, together with the separate RMF in its hibernation system (Fig. 5B and S8A). To examine how these HPF regions influence translation in a different bacterial translation environment, we tested the corresponding HPF variants in an *E. coli* cell-free translation system (Fig. 5).

**Figure 5.**
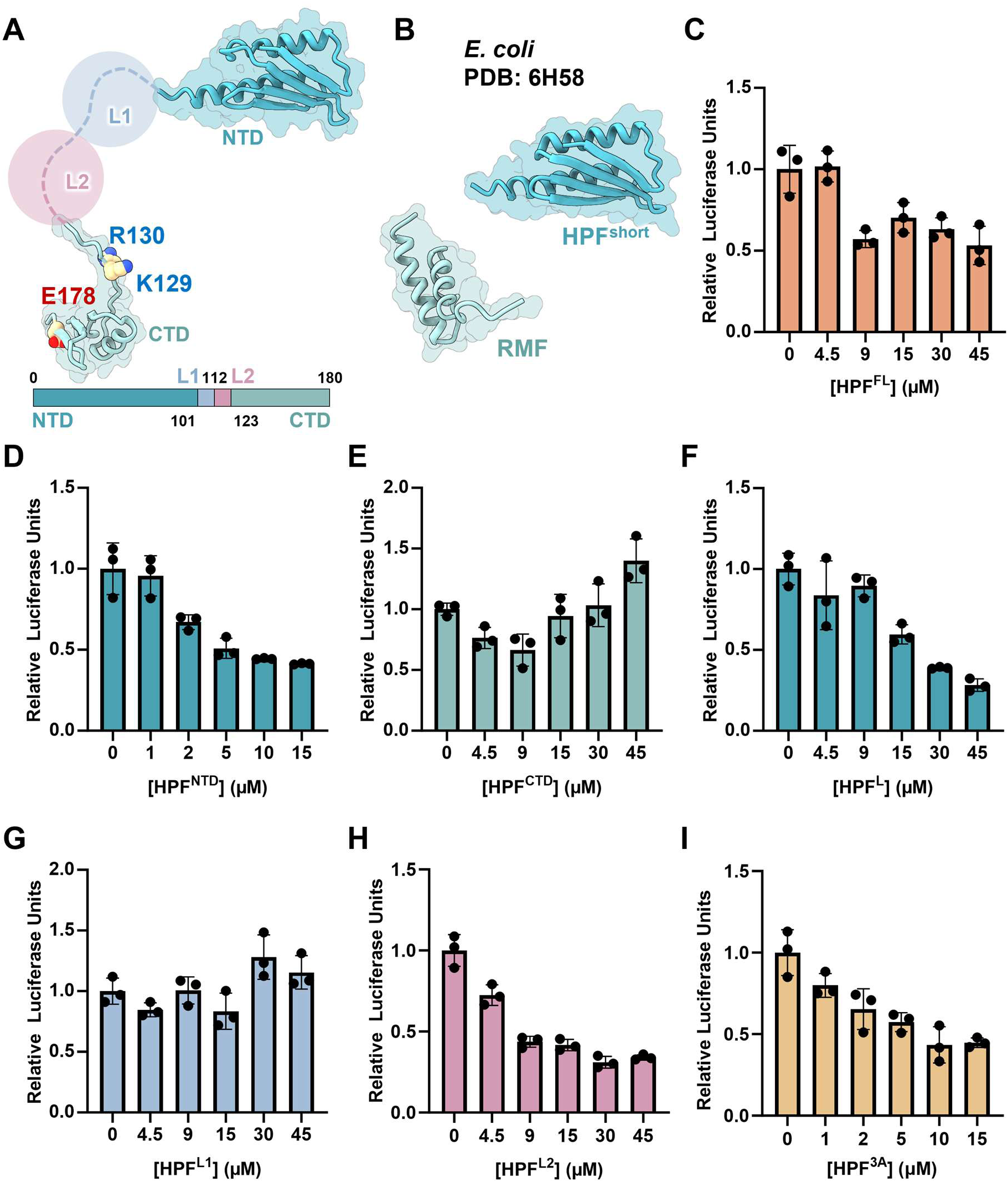
Analysis of HPF variants in *E. coli*. (**A**) Structural model of *Bacillus* HPF, with flexible Linker regions L1 and L2 shown as dashed lines due to lack of ordered density. Key residues K129, R130, and E178 forming the electrostatic pocket in the CTD are shown as spheres. The bottom schematic indicates the domain organization of long-form HPF. **(B)** Structural representation of the *E. coli* hibernation system (PDB: 6H58), showing short-form HPF and RMF (Beckert *et al*, 2018). (**C-I)** IVT assays evaluating the heterologous translational repression by *Bacillus* HPF variants using a NanoLuciferase reporter in the *E. coli* cell-free translation system. Data are presented as mean ± s.d. (n = 3).

We first compared full-length HPF with the isolated NTD and CTD (Fig. 5C-E). HPF^FL^ produced moderate translation inhibition (Fig. 5C) whereas the isolated HPF^NTD^ exhibited strong dose-dependent inhibition and HPF^CTD^ showed little inhibitory activity (Fig. 5C-E). Notably, the HPF^NTD^ displayed relatively weak activity in the native *Bacillus* system (Fig. 2K), suggesting that the contribution of the isolated NTD to translation inhibition can vary with the translational context. The Linker variants further revealed differential effects on translational repression, with HPF^L2^ showing stronger repression than HPF^L^, whereas HPF^L1^ exhibited markedly weaker activity (Fig. 5F-H). We then examined the charged CTD surface positioned near the aSD-containing 16S rRNA 3′ terminus using the HPF³ᴬ mutant. HPF³ᴬ exhibited stronger translational repression than HPF^FL^ across the tested concentrations (Fig. 5I). These results suggest that the charged CTD surface contributes to translational regulation beyond its role in CTD-mediated ribosome dimerization.

Based on the structural and functional analyses, we propose a model for how these modules are integrated within the long-form HPF-100S complex (Fig. 6). The CTD forms a hydrophobic inter-ribosomal interface that promotes 100S assembly, while the Linker connects the NTD and CTD and is positioned near the 16S rRNA 3′ terminus containing the aSD sequence. The weaker inhibitory activity of HPF^L1^ in both translation systems suggests that Linker1 may influence the functional activity or positioning of the NTD, although the underlying mechanism remains unclear. The charged CTD surface adjacent to the rRNA 3’ region, together with the Linker, may provide additional regulatory inputs that influence HPF-mediated translational repression. In this model, the CTD, Linker, and NTD form an integrated regulatory structure that coordinates 100S assembly with translational repression.

**Figure 6.**
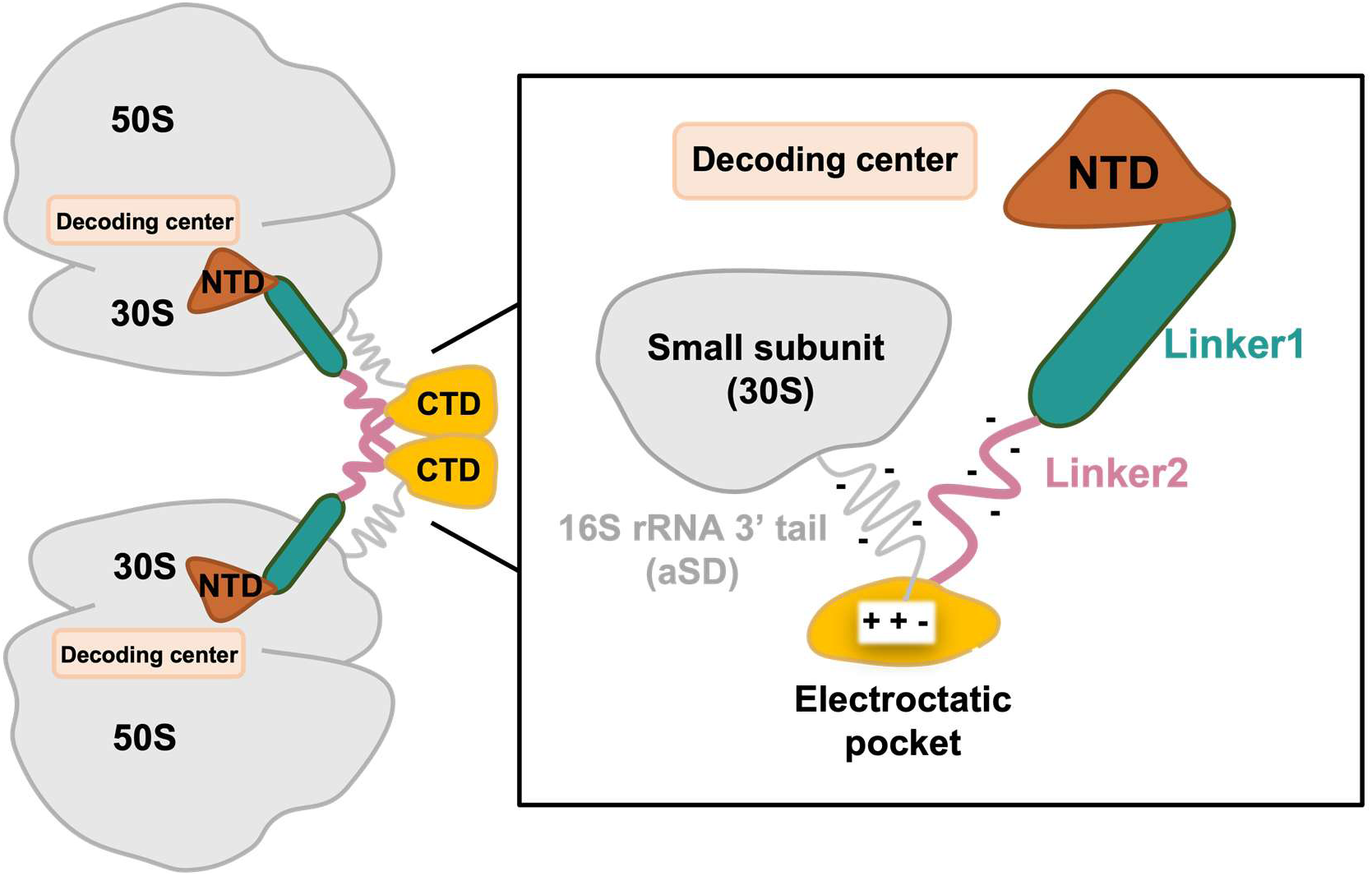
Proposed working model for long-form HPF in ribosome hibernation. Schematic showing the modular domain architecture of long-form HPF in coordinating 100S disome assembly and translational repression. The NTD (brown) inserts into the decoding center of the 30S subunit to block translation. The CTD (yellow) drives inter-ribosomal dimerization. Linker 1 (green) serves as a force-transmitting tether, while Linker 2 (pink) and the rRNA-facing electrostatic pocket of the CTD modulate conformational tension with the flexible 16S rRNA 3′ containing the aSD sequence.

## Discussion

Ribosome hibernation enables bacteria to preserve translational capacity during prolonged environmental stress, yet how hibernating complexes achieve stable translational arrest while maintaining the capacity for rapid reactivation remains poorly understood. In Gammaproteobacteria, such as *E. coli*, short-form HPF cooperates with RMF to regulate decoding-center accessibility and aSD sequence sequestration (Wada *et al*, 1990; Beckert *et al*, 2018; Yoshida *et al*, 2021; Polikanov *et al*, 2012; Kato *et al*, 2010), whereas Firmicutes employ a single long-form HPF protein to establish ribosome dormancy (Beckert *et al*, 2017; Khusainov *et al*, 2017; Matzov *et al*, 2017; Kumar *et al*, 2024; Chan *et al*, 2025; Franken *et al*, 2017). Here, our cryo-EM structure and biochemical analyses of a native deep-sea *Bacillus* 100S disome reveal that long-form HPF integrates functions that are distributed between the two-factor *e*HPF/RMF system. By combining decoding-center occlusion through the NTD, CTD-mediated disome dimerization, and Linker-CTD-associated organization positioning near the 16S rRNA 3′ terminus, long-form HPF provides a modular mechanism for coupling ribosome dimerization with translational repression.

One insight from our native disome reconstruction is that the hibernating ribosome exists as a dynamic conformational ensemble rather than a uniformly locked inactive particle. Earlier studies of long-form HPF-containing 100S particles, including the hibernating ribosomes from *Staphylococcus aureus* and *Thermus thermophilus* (Khusainov *et al*, 2017; Flygaard *et al*, 2018), indicated that dormant ribosomes can adopt both rotated and unrotated conformations, suggesting that the hibernating state is not structurally static. Our structure extends these observations by showing that conformational heterogeneity is retained within a native HPF-bound disome population, with most particles adopting a symmetrical non-rotated state while a smaller population samples an asymmetric partially rotated state. Such conformational flexibility may provide a structural basis for maintaining a repressed ribosome state while permitting structural transitions within the hibernating ensemble. Thus, the 100S disome represents a structurally heterogeneous hibernation state rather than a uniformly locked inactive particle.

Our functional dissection of HPF reveals an unexpected modular organization underlying ribosome control. CTD-mediated inter-ribosomal interactions provide the major structural interface for 100S assembly, whereas the NTD and Linker-containing region contribute to translational regulation through distinct mechanisms. The strong inhibitory activity of the isolated NTD in the *E. coli* translation system further supports the NTD as a key module for translational repression, while the differential effects of Linker substitutions indicate that the intervening region modulates this activity rather than serving simply as a passive connection between the NTD and CTD. In particular, the weaker activity of Linker 1 substitutions across translation systems raises the possibility that this segment influences the functional state of the NTD-containing region. Importantly, mutation of charged residues within the CTD-associated electrostatic surface does not impair disome formation, despite enhancing translational repression *in vitro*. This finding suggests that the CTD contributes to HPF function beyond its role in inter-ribosomal association, and that the charged surface near the aSD-containing 16S rRNA 3′ terminus may modulate the functional state of the HPF-ribosome complex. One possibility is that alterations in this electrostatic surface influence the local environment of the HPF-ribosome complex, thereby affecting the efficiency of NTD-mediated translation inhibition. Although the precise molecular basis requires further investigation, the combined structural and functional data support a model in which the NTD, Linker, and CTD-associated surfaces make distinct but interconnected contributions to ribosome hibernation.

Together, our study expands the current understanding of bacterial ribosome hibernation by revealing how conformational flexibility and modular HPF functions are combined within the 100S disome. The long-form HPF provides a single-factor framework for coordinating ribosome storage, translational repression, and regulatory flexibility. Through functional specialization of its domains and conformational heterogeneity of the 100S particle, HPF contributes to a regulated dormant state that enables bacteria to conserve energy during stress while maintaining the potential for subsequent translational reactivation. These findings highlight ribosome hibernation as a regulated and dynamic state rather than a simple endpoint of translational shutdown.

## Methods

### Isolation, cultivation, and characterization of *Bacillus* sp. DY0427

Deep-sea sediment samples used for bacteria isolation in this study were collected from a hydrothermal vent (26.012° N, 13.856° W) at a depth of 4,346 m. A basal medium was prepared using filtered seawater (pH 7.0) supplemented with 1.0 g/L yeast extract, 1.0 g/L peptone, 1.0 g/L NaNO_3_, 1.0 g/L NH_4_Cl, 1.0 g/L NaHCO_3_, 1.0 g/L CH_3_COONa, 0.5 g/L KH_2_PO_4_, 0.2 g/L MgSO_4_.7H_2_O, 0.7 g/L cysteine hydrochloride, 500 µL/L 0.1% (w/v) resazurin. Enrichment was carried out under a 100% N_2_ atmosphere at 28 °C for one month. For isolation, the basal medium was solidified with 15 g/L agar and evenly coated onto the inner wall of Hungate tubes to form a thin layer. After enrichment, 50 µL of the culture was transferred anaerobically into a roll tube and spread onto the medium layer. The tubes were incubated anaerobically at 28 °C for three days. Single colonies were picked using sterilized bamboo sticks and subsequently cultured in 15 mL Hungate tubes containing 10 mL of basal medium at 28 °C for three days. These microbial species were identified by PCR. A deep-sea *Bacillus* strain (designated DY0427) was isolated and purified by repeated application of the Hungate roll-tube method. Purity was routinely confirmed by transmission electron microscopy (TEM) and partial 16S rRNA gene sequencing.

The morphology of *Bacillus* sp. DY0427 was examined by transmission electron microscopy. A 10 mL culture was harvested by centrifugation at 5,000 g for 10 min. The resulting pellet was washed three times with phosphate-buffered saline (PBS; 137 mM NaCl, 2.7 mM KCl, 10 mM Na_2_HPO_4_ and 1.8 mM KH_2_PO_4,_ pH 7.4) and resuspended in 20 μL of PBS. Carbon film-coated copper grids were immersed in the cell suspension for 20 min, and the prepared samples were examined using an HT7700 transmission electron microscope (Hitachi, Japan).

### Isolation and purification of *Bacillus* ribosomes

To isolate *Bacillus* ribosomes, the cell pellet was resuspended in lysis buffer (20 mM HEPES-KOH pH 7.5, 10 mM Mg(OAc)_2_, 100 mM NH_4_Cl, 2 mM DTT). Cells were lysed using a high-pressure homogenizer (1290 bar, 4 passes), and the lysate was centrifuged at 15,000 g for 10 min at 4 °C to remove cell debris. The supernatant was loaded onto a 0.8 M sucrose cushion and centrifuged at 603,000 *g* for 50 min at 4 °C using an MLA-130 rotor. The pellet was resuspended in 500 μL of lysis buffer, and the crude ribosome sample was loaded onto a 10% - 40% sucrose density gradient prepared in lysis buffer. The sucrose gradient was transferred to a Beckman-Coulter SW41 rotor and centrifuged at 200,000 g for 2.5 h. After centrifugation, the gradient was collected using a fractionation system. Gradient samples corresponding to different peaks were collected respectively. The fraction corresponding to the 100S peak was buffer-exchanged into grid buffer (20 mM HEPES-KOH pH 7.5, 10 mM Mg(OAc)_2_, 30 mM KCl, 1 mM DTT), and was immediately subjected to cryo-EM grid freezing.

### Cryo-EM sample preparation

The concentrations of the *Bacillus* ribosomes used for cryo-EM grid preparation were 170 nM. Approximately 3.5 μL aliquots were applied to a plasma-cleaned holey carbon grid (R1.2/1.3, 300 mesh, Quantifoil) for 1 min, on which a homemade continuous carbon film was precoated. Grids were blotted for 2.5 s under 100% humidity at 4 °C and plunge-frozen in liquid ethane using Vitrobot Mark IV (Thermo Fisher Scientific).

### Cryo-EM data collection and processing

Cryo-EM micrographs of all samples were collected on a Titan Krios transmission electron microscope (Thermo Fisher Scientific) operated at 300 kV using a K3 camera and a slit width of 20 eV on a GIF quantum energy filter (Gatan). Movies were recorded in super resolution mode at a nominal magnification of 105,000×, corresponding to a calibrated physical pixel size of 0.41 Å per pixel. A total electron dose of 50 electrons per Å² was fractionated over 32 frames with a total exposure time of 2.3 s. Data were acquired using EPU software with a defocus ranging from 1.0 to 2.0 μm. In total, 6,825 movies were collected.

Datasets were processed using cryoSPARC v.4.5.3 (Punjani *et al*, 2017). Movie stacks were motion-corrected using the patch motion correction tool with 2× binning (bin2). CTF parameters were estimated using the patch CTF estimation tool. Particles were initially picked using template-free blob picker tool, extracted, and subjected to 2D classification. Class averages displaying well-resolved 70S ribosome features were selected and used as templates for template-based particle picking. The newly picked particles underwent multiple rounds of 2D classification to discard junk particles and contaminants. The cleaned particle set was then used for ab initio reconstruction, and the resulting volumes were further refined by heterogeneous refinement; poor-quality classes were discarded at this stage. The retained particles were re-extracted using an enlarged box size sufficient to accommodate the 100S ribosome. These particles were then subjected to additional 2D classification, and a final curated stack of high-quality particles was selected for homogeneous refinement. After homogeneous refinement of the 100S ribosome particles, one monomer displayed pronounced conformational flexibility and poor local resolution. To address this heterogeneity, focused 3D classification was performed, yielding a subset in which both monomers were well-resolved. This stable subset was further subjected to 3D classification, which separated the particles into two distinct rotational states: a non-rotate/non-rotate state and a rotate/non-rotate state. Each state was subsequently refined by homogeneous refinement to generate the final density maps. To enhance local resolution at key functional sites, spherical masks were generated using the “relion_mask_create” tool in RELION 4.0 (Kimanius *et al*, 2021; Scheres, 2012) and applied during local focused refinement of each class (Scheres & Chen, 2012). Map resolutions are reported based on the FSC = 0.143 criterion.

### Model building and refinement

Genome sequencing data from *Bacillus* were used to obtain rRNA and protein sequences. Initial ribosomal RNA (rRNA) models were predicted by AlphaFold3 (Abramson *et al*, 2024), and initial protein models were built using CryoAtom (Su *et al*, 2026). The model was inspected for completeness, and regions with missing or incorrect sequences were manually rebuilt. The final models were further refined using Coot v0.9.8.2 (Emsley & Cowtan, 2004; Emsley *et al*, 2010) and Phenix.real_space_refine v1.21.2 (Afonine *et al*, 2018; Liebschner *et al*, 2019). Figures were prepared with ChimeraX v1.8 (Meng *et al*, 2023) and Graphpad Prism 10.1.2.

### Recombinant protein expression and purification

The gene encoding the HPF protein from the *Bacillus* was amplified from genomic DNA by PCR and cloned into the pET-28a(+) prokaryotic expression vector with an N-terminal 6× His tag. Correct insertion was verified by double digestion with *Nco*I and *Xho*I restriction enzymes. HPF protein was overexpressed in *E. coli* BL21(*DE3*). Cells were cultured at 37 °C until the OD_600_ reached 0.8, and then 0.5 mM IPTG was added to induce HPF overexpression. The culture was further incubated at 25 °C and 180 rpm for 16 h, after which the cell pellet was collected by centrifugation at 3,500 *g* for 30 min.

The cell pellet was resuspended in lysis buffer (20 mM HEPES-KOH pH 8.0, 20 mM KCl, 20 mM MgCl₂, 500 mM NaCl, 10% glycerol, 1 mM PMSF) and lysed by four passes through a high-pressure homogenizer at 800 bar. The lysate was centrifuged at 20,000 g for 30 min at 4 °C, and the supernatant was collected and incubated with Ni-NTA affinity resin (Cytiva) for 1-2 h at 4 °C on a rotator.

After incubation, the resin was washed three times with wash buffer (20 mM HEPES-KOH pH 8.0, 20 mM KCl, 20 mM MgCl₂, 500 mM NaCl, 10% glycerol, 30 mM imidazole) to remove non-specifically bound proteins. The protein was then eluted with 20 mL of elution buffer (20 mM HEPES-KOH pH 8.0, 20 mM KCl, 20 mM MgCl₂, 500 mM NaCl, 10% glycerol, 300 mM imidazole), and 1 mL fractions were collected. Fractions containing the target protein were pooled and concentrated using an ultrafiltration device with a 10 kDa molecular weight cutoff (Millipore). Protein purity was verified by SDS-PAGE. The pooled HPF protein fractions were further concentrated by ultrafiltration, and the protein concentration was determined by measuring the absorbance at 280 nm. The final protein sample, at a concentration of approximately 6 mg/mL, was aliquoted and stored at −80 °C.

### Western blotting

For Western blot analysis, samples from the binding reaction in which ribosomes were incubated with HPF protein at a molar ratio of 1:1 were mixed with SDS loading buffer (containing 5% β-mercaptoethanol) to a final concentration of 1×, and denatured in a boiling water bath for 5 min. The protein samples were then separated by 15% SDS-polyacrylamide gel electrophoresis (SDS-PAGE) using a Tris-glycine running buffer system (25 mM Tris, 192 mM glycine, 0.1% SDS). After electrophoresis, the separated proteins were transferred onto a 0.22 μm PVDF membrane (Millipore) by the wet transfer method. The transfer was carried out at a constant current of 200 mA for 90 min in transfer buffer composed of 25 mM Tris, 192 mM glycine, and 20% methanol.

Following the transfer, the PVDF membrane was blocked with 5% non-fat dry milk dissolved in TBST for 1 h at room temperature. The membrane was then washed three times with TBST buffer (20 mM Tris-HCl, pH 7.5, 150 mM NaCl, 0.1% Tween 20), 5 min for each wash. After washing, the membrane was incubated with the corresponding primary antibody overnight at 4 °C on a shaker. The primary antibody used was an anti-His tag mouse monoclonal antibody (Zenbio, catalog number: 230001) diluted at a ratio of 1:3000. After the primary antibody incubation, the membrane was washed three times with TBST buffer, 10 min for each wash, and subsequently incubated with the corresponding HRP-conjugated secondary antibody, a goat anti-mouse IgG (Zenbio, catalog number: 511103), at a dilution ratio of 1:10000 for 1 h at room temperature. The membrane was then washed again three times with TBST buffer, 10 min per wash. Chemiluminescent detection was performed using an ECL chemiluminescence detection kit (Vazyme, catalog number: E422-01), and the signals were captured with an iBright imaging system.

### *In vitro* translation assay

The purified proteins HPF^FL^, HPF^NTD^, HPF^CTD^, HPF^L1^, HPF^L2^, HPF^L^, and HPF^3A^ were tested for inhibitory activity in a cell-free translation system (Köhrer *et al*, 1996; Hall *et al*, 2012). The *Bacillus* cell lysate was prepared as follows: cells were collected by centrifugation at 3,500 g for 15 min, washed 2-3 times with 1× PBS, and resuspended in lysis buffer (10 mM HEPES-KOH pH 7.5, 14 mM Mg(OAc)_2_, 60 mM KOAc) at a ratio of 1 g/mL. The cells were lysed by a high-pressure homogenizer (1290 bar, 4 passes), and the supernatant was clarified by centrifugation at 15,000 g for 30 min. A 10 μL in vitro translation reaction mixture was assembled as follows: 3 µL cell lysate, 1 µL salt mix, 1 µL NTP mix, 1 µL energy mix, 1 µL amino acid mix, 0.17 µg tRNA, 0.93 µg T7 RNA polymerase, 68 ng DNA, and 2 μL of protein or storage buffer. The reaction was incubated at 30 °C for 30 min, and luciferase activity was measured using a microplate luminometer (BERTHOLD Centro XS3 LB 960). To control for batch-to-batch variation, lysate and mRNA aliquots from the same preparation batch were used for comparative assays within each experimental series.

### Ribosome binding assay

Cell lysate from *Bacillus* cells was centrifuged through a 1 M high-salt sucrose cushion buffer (20 mM HEPES-KOH, 1 M KOAc, 10 mM Mg(OAc)₂, 1 mM DTT, 1 M sucrose) to obtain vacant ribosomes. The ribosomes were resuspended in resuspension buffer (10 mM HEPES-KOH, 100 mM KOAc, 10 mM Mg(OAc)₂, 10 mM NH₄Cl, 0.25 mM EDTA, 1 mM DTT) and diluted to 2 μM. Purified HPF^FL^, HPF^NTD^, HPF^CTD^, HPF^L1^, HPF^L2^, HPF^L^, and HPF^3A^ proteins (6×His-tagged) at 2 μM and 20 μM were individually incubated with the ribosomes at 4 °C to form complexes. The reaction mixtures were centrifuged through a 1 M sucrose cushion buffer (containing 100 mM and 500 mM K⁺) (Beckman Coulter MLA130 rotor, 603,000 g, 4 °C, 50 min) to separate ribosomes and associated factors. The ribosomal pellets were resuspended in resuspension buffer and analyzed by Western blotting (WB). Quantitative agarose gel electrophoresis was performed using a 1.5% (w/v) agarose gel in 1× TAE buffer at 180 V for 25 min. Gels were imaged using an iBright imaging system.

### Sucrose density gradient separation of polysomes formed by protein incubation

Cell lysate from *Bacillus* cells was centrifuged through a 1 M high-salt sucrose cushion buffer (20 mM HEPES-KOH, 1 M KOAc, 10 mM Mg(OAc)₂, 1 mM DTT, 1 M sucrose) to obtain vacant ribosomes. After resuspending the pellet in resuspension buffer, a 10%-40% continuous sucrose density gradient (with ionic conditions identical to the resuspension buffer) was prepared using a gradient former. The gradient was centrifuged at 200,000 g for 2.5 h at 4 °C to separate pure 70S ribosomes. The monomer peak was collected, and ribosomes were pelleted by centrifugation at 603,000 g for 50 min. The ribosomes were resuspended in resuspension buffer, diluted to 1 μM, aliquoted, and stored at −80 °C.

Ribosomes purified by high-salt sucrose cushion and continuous density gradient centrifugation and stored at −80 °C were incubated with 1 μM each of purified HPF^FL^, HPF^NTD^, HPF^CTD^, HPF^L1^, HPF^L2^, HPF^L^, and HPF^3A^ proteins at 4 °C to form complexes. The reaction mixtures were subjected to 15%-30% continuous sucrose density gradient centrifugation to separate ribosomal subunits, monomers, and dimers, and the change in dimer content was observed.

## Data availability

The cryo-EM maps and atomic coordinates generated in this study have been deposited in the Electron Microscopy Data Bank and Protein Data Bank under accession numbers EMD-83096 and 44YC, respectively.

## Author contributions

W.L. and C.S. conceived the study. R.Z. provided *Bacillus* sp. DY0427. Y.B. and C.Z. performed the biochemistry experiments including the *in vitro* translation, ribosome binding assay, sucrose gradients and cryo-EM samples preparations. The cryo-EM data were collected at the Biomedical Research Center for Structural Analysis of the Shandong University. M.W., L.Q., C.Y. and L.Z. performed the cryo-EM data collection, image processing, model building, and structural analysis. M.W., C.Y., L.Z., Y.B., C.Z. prepared the figures. W.L. supervised the project and wrote the initial draft of the manuscript. All authors analyzed the data and edited the paper.

## Disclosure and competing interest statement

The authors declare no competing interests.

## Acknowledgements

The cryo-EM data were collected at the Biomedical Research Center for Structural Analysis of the Shandong University. This work was supported by grants from the National Natural Science Foundation of China (32371351 and 32171291 to W.L., 82302100 to M.W.), Shandong Basic Research Special Zone (Basic Medical Sciences) (TQ042025005 to W.L.), Cutting Edge Development Fund of Advanced Medical Research Institute (GYY2023QY01 to W.L.) and Shandong Excellent Young Scientists Fund Program (2022HWYQ-025 to W.L.).

